# Micafungin-induced Cell Wall Damage Stimulates Microcycle Conidiation in *Aspergillus nidulans*

**DOI:** 10.1101/2021.05.19.444817

**Authors:** Samantha Reese, Cynthia Chelius, Wayne Riekhof, Mark R. Marten, Steven D. Harris

## Abstract

Fungal cell wall receptors relay messages about the state of the cell wall to the nucleus through the Cell Wall Integrity Signaling (CWIS) pathway. The ultimate role of the CWIS pathway is to coordinate repair of cell wall damage and to restore normal hyphal growth. Echinocandins such as micafungin represent a class of antifungals that trigger cell wall damage by affecting synthesis of β-glucans, filamentous fungi’s response to these antifungals are fundamentally unknown. To obtain a better understanding of the dynamics of the CWIS response and its multiple effects, we have coupled dynamic transcriptome analysis with morphological studies of *Aspergillus nidulans* hyphae responding to micafungin. Our results reveal that expression of the master regulator of asexual development, BrlA, is induced by micafungin exposure. Further study showed that micafungin elicits microcycle conidiation in a BrlA-dependent manner, and that this response is abolished in the absence of MpkA. Our results suggest that microcycle conidiation may be a general response to cell wall perturbation which in some cases would enable fungi to tolerate or survive otherwise lethal damage.

## 1. Introduction

The cell wall is an important feature of filamentous fungi, where it is responsible for the protection of hyphal integrity and the maintenance of hyphal morphology [1]. In fungi, damage to the cell wall poses a serious threat to cell viability. To mitigate this possibility, fungi employ cell surface sensors that detect damage to the cell wall and activate a suite of mechanisms that repair the damage and enable the resumption of normal growth [1]. This response is known as the Cell Wall Integrity Signaling (CWIS) pathway [2]. This pathway is found in many fungal species and as key signaling features includes the Rho1 GTPase, protein kinase C, and the CWIS MAP kinase cascade [2]. In *Aspergillus nidulans*, the pathway is conserved and terminates with the MAP kinase MpkA [3].

There are many perturbations that can trigger the CWIS pathway in fungi including antifungals detected in the environment. Micafungin, an echinocandin antifungal drug, has been shown to activate the CWIS pathway. It acts by inhibiting the activity of 1,3-beta glucan synthase, thereby depleting β-1,3 glucan from the cell wall and compromising hyphal integrity [4]. This leads to hyphal tip bursting and a decrease in internal hydrostatic pressure of the hyphae [5]. The CWIS response to micafungin is well understood in yeast, but is less studied in filamentous fungi such as *A. nidulans*. Notably, the transcriptional output of the CWIS pathway differs significantly in *A. nidulans* than in the yeast *Saccharomyces cerevisiae* [3]. To better understand the dynamic nature of the CWIS in *A. nidulans*, we have employed a “multi-‘omics” approach aimed at characterizing key effectors and outputs of micafungin perturbation [6].

The asexual life cycle of filamentous fungi is important for maintaining species viability in the face of adverse environmental conditions [7]. The formation of spores via asexual means is typically a well-regulated global response to both external and internal stimuli [8]. However, under some circumstances, such as specific environmental triggers, the normal regulatory and developmental processes that underlie asexual sporulation can be short-circuited to allow rapid production of spores [9]. The shortened cycle for the production of conidia by newly germinated spores without any further hyphal growth is termed microcycle conidiation [10]. At this time, the overall benefits of this process and the extent to which fungi use it as a short-term survival strategy remain unclear. Microcycle conidiation has been observed across many ascomycetes, including members of the genera *Aspergillus, Fusarium*, and *Metarhizium* [8, 9, 11]. In these fungi, the genetic pathways that regulate microcycle conidiation and coordinate its occurrence relative to environmental conditions are poorly understood in limited detail.

The objective of this study was to gain a deeper understanding of the filamentous fungal response to cell wall damage by exposing the model fungus *A. nidulans* response to micafungin exposure as a paradigm. Upon exposure of growing hyphae to micafungin, samples were taken over a time course and subjected to both phosphoproteomic and transcriptomic analyses [6]. Here, we describe the impacts of exposure to micafungin on patterns of gene expression in both wildtype and Δ*mpkA* mutants. Our results suggest that the response to cell wall damage is attenuated in Δ*mpkA* mutants, and reveal specific classes of genes whose expression is dependent upon MpkA. Notably, this includes the master regulator of conidiation, *brlA*. Indeed, we show that BrlA-dependent microcycle conidiation is triggered by exposure to micafungin, thereby raising the possibility that this might be a critical feature of the response to the class of anti-fungal drugs.

## 2. Materials and Methods

### 2.1 Media and Growth Conditions

All strains used in this study are described in **Supplementary Table 1**. Standard media and growth conditions were employed [12, 13]. Media included Yeast Extract-Glucose-Vitamin (YGV; 0.5% yeast extract, 1% glucose, 0.1% vitamins [13], Malt Extract-Glucose (MAG; 2% malt extract, 2% glucose, 0.2% peptone, 0.1% trace elements, 0.1% vitamins [13].

### 2.2 Global Analysis Strain Generation, Growth Conditions, RNA Extraction, and RNA Sequencing

The generation of all strains, growth conditions for RNA-Sequencing samples, RNA extraction and RNA Sequencing pipeline can be found in our previous paper [6]. NCBI Sequence Read Archive Accessions can be found in (**Supplementary Table 2**).

### 2.3 qRT-PCR for RNA-Sequencing Verification

Wildtype and Δ*mpkA* hyphae grown in YGV for 11 hours were exposed to 1ul of micafungin per 1ml growth media for 75 minutes [12]. Control hyphae were left untreated [12]. The hyphae were frozen with liquid nitrogen, and RNA was then extracted and purified. A cDNA library was generated using the cDNA Library Kit from Thermo Fisher Scientific (catalog A48571) and the Reverse transcription PCR kit from Millipore Sigma (Product No. HSRT100), with specific primers were designed for each target gene (**Supplementary Table 3**). The samples were analyzed using Bio-Rad CFX Manager 3.1. There were three biological replicates, and the fold-change was determined by comparing the treated and untreated samples using the ΔΔCt method [14]. Histone *H2B* was used as a control for qRT-PCR normalization.

### 2.4 Microscopy

Images were collected using either an Olympus BX51 microscope with a reflected fluorescence system fitted with a Photometrics CoolSnap HQ camera coupled to Lambda B10 Smart Shutter control system (Sutton Instruments), or an EVOS FL microscope (Advanced Microscopy Group). Images were initially processed using MetaMorph software (Molecular Devices).

### 2.5 Micafungin Concentration Experiments

Wildtype and Δ*mpkA* hyphae were inoculated and grown as mentioned above on cover slips in 60mm X 15mm petri dishes for 11 hours in 25ml of YGV media. At this time, micafungin (0.0ng/ml, 0.01ng/ml, 0.1ng/ml, 1.0ng/ml) was added and hyphae were grown for an additional three hours before viewing.

### 2.6 Microcycle conidiation Experiment with Wildtype and ΔmpkA

Wildtype and Δ*mpkA* hyphae were inoculated and grown as mentioned above with the micafungin dosage of 0.1ng/ml in a total of 25ml of YGV media. Hyphae were grown an additional nine hours post antifungal addition. Calcofluor white (Catolog number 18909-100ML-F Sigma) was added to cover slips before viewing at 0.1ng/ml to enhance visualization of spore cell wall.

### 2.7 Up-regulation of brlA Experiments

The *alcA::brlA* strain constructed by Adams et al. (1988) was grown on cover slips in YGV for 11 hours, then the cover slips were shifted into MNV 100mN L-Threonine to induce the *alcA* promoter [15,16]. After an additional three hours of growth in MNV 100mN L-Threonine media, calcofluor white was added to cover slips at 0.1ng/ml to enhance visualization of spore cell wall [12].

## 3. Results

### 3.1 Global Transcript Expression

To better understand the dynamic response of the CWIS pathway to cell wall damage, hyphae exposed to micafungin were harvested at 0, 5, 10, 15, 20, 25, 30, 40, 50, 60, 75, 90, 120 minutes, and subjected to transcriptomic analysis as described in Chelius et al. (2020) [6]. The zero minute time point had no micafungin exposure and served as our untreated control for transcript expression. Significantly expressed genes (p-value ≤ 0.1) were identified following analysis using DESeq2 in RStudio [17]. The wildtype strain showed an exponential increase of significantly expressed genes beginning at 25 minutes through to the last time point with the total of 1614 genes showing altered expression (**Figure 1A and Supplementary Table 4**). The Δ*mpkA* strain showed a linear increase in significantly expressed genes from beginning at 30 minutes through to the last time point with a total of 964 differentially expressed genes (**Figure 1A and Supplementary Table 5**).

**Figure 1:**
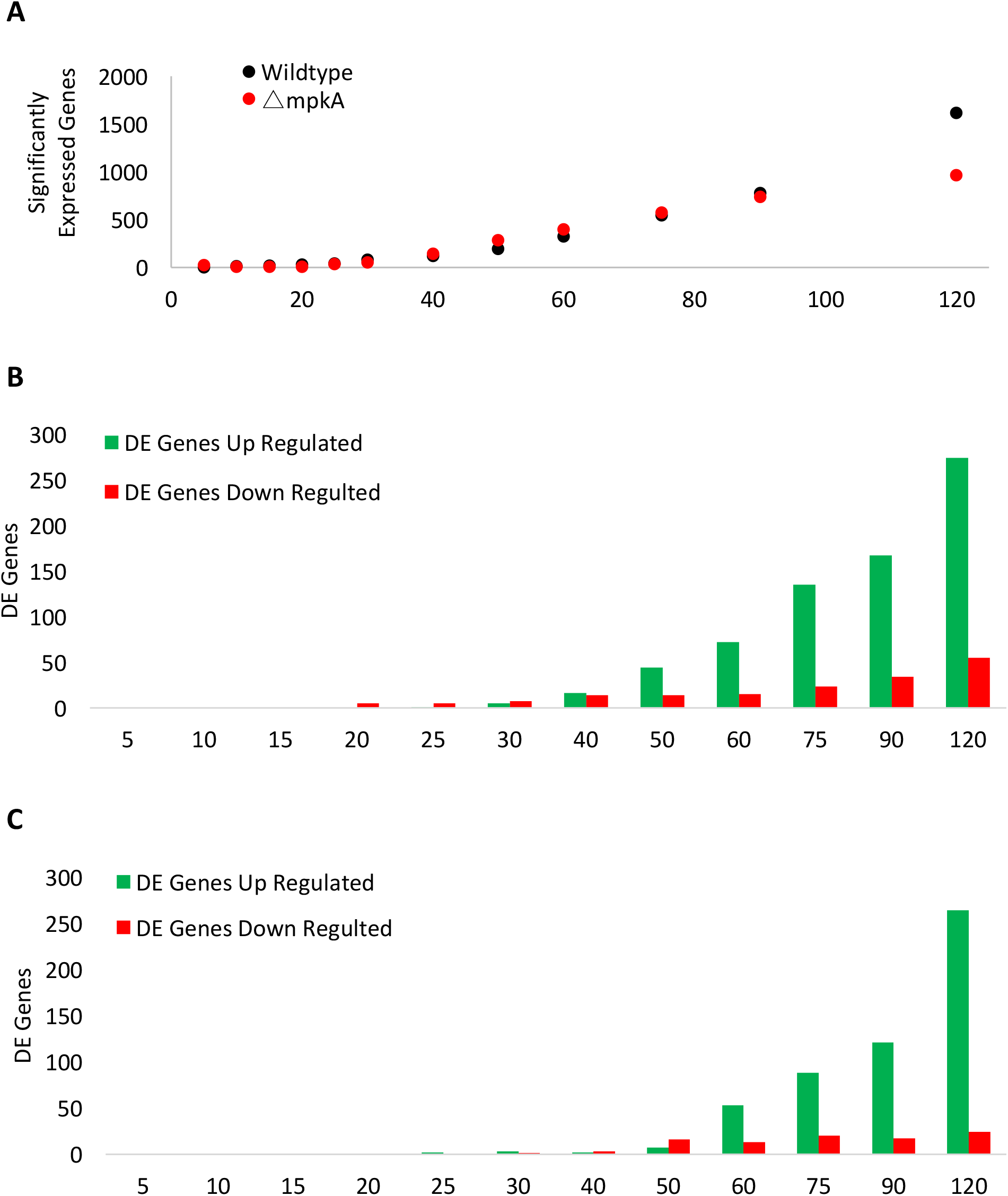
Global Gene Expression. **A)** Wild type showed exponential increase in significantly expressed genes post micafungin exposure. Δ*mpkA* showed linear increase in significantly expressed genes post micafungin exposure. Data points for both up-regulated and down-regulated genes. (Significantly expressed genes = p-value < 0.1). Three biological replicates. **B)** Wildtype DE genes. The first DE gene in wild type was at the 15 minutes. **C)** Δ*mpkA* DE genes. The first DE gene in Δ*mpkA* was at 25 minutes. (DE genes = p-vale < 0.1, Log2Fold change (+/-) 2.0) Three biological replicates. X-axis is minutes.

There was a linear increase of both up and down-regulated Differently Expressed (DE) genes (adjusted p-value ≤ 0.1 and log2fold (+/-) 2.0) in both wildtype and Δ*mpkA* starting at the 30-minute time point (**Figure 1B, Figure 1C**, and **Supplementary Table 6**). There also were more up-regulated DE genes in both wildtype and Δ*mpkA* strains. In wildtype, 274 DE genes were up-regulated and 55 DE genes down-regulated, whereas in Δ*mpkA* 264 DE genes were up-regulated and 24 DE down-regulated (**Figure 1B, Figure 1C**, and **Supplementary Table 6**). Some of the DE genes were up-regulated in both wildtype and Δ*mpkA* (**Supplementary Figure 1)**. The first shared DE genes were AN1199, AN3888, and AN8342 at time point 50 minutes. The number of shared DE genes increased throughout the time course, such that at the last time point there were eighty-five shared genes between wildtype and Δ*mpkA*. These genes seemingly defined a component of the CWIS that is independent of MpkA.

### 3.2 Gene Cluster Regulation

Analysis of DE genes in both wildtype and the Δ*mpkA* strain at each time point revealed that a number of apparent gene clusters (defined as three or more genes with consecutive ANID numbers) were differentially regulated upon exposure to micafungin (**Supplementary Table 7)**. For example, as early as 40-minutes post micafungin exposure, expression of the AN5269-5273 cluster was up-regulated in wildtype whereas that of the AN7952-7955 cluster was up-regulated in the Δ*mpkA* strain. A total of 19 presumptive gene clusters exhibited dynamic gene expression, with the largest number at the final time point. These clusters could be sorted into three groups whose expression was affected by micafungin; (i) MpkA-dependent micafungin induced, (ii) MpkA-independent micafungin-induced, and (iii) MpkA-dependent micafungin-repressed. However, the largest group consisted of DE clusters that were dependent on MpkA for expression but were not affected by exposure to micafungin. Some of these clusters were also identified in our earlier study of MpkA-dependent gene expression in the absence of cell wall stress [18]. With some exceptions (Andersen et al., 2013), the precise function of the apparent clusters within the DE set of gene is unknown, but this observation highlights the complexity of the CWIS and the presence of both MpkA-dependent and -independent components [10].

### 3.3 GO Term Analysis

The distribution of DE genes amongst Gene Ontology (GO) categories were analyzed using the *Aspergillus* Database [19, 20]. In wildtype the up-regulated GO term categories showed a shift in gene expression starting at the 60-minute time point (**Figure 2A** and **Supplementary Table 8**). From the 25-minute time point to the 60-minute time point, the data revealed a response to micafungin that featured increases in oxidoreductase activity genes including; AN7708 (Log2Fold 2.25), AN3043 (Log2Fold 2.66), AN9375 (Log2Fold 2.71), *uaZ* (Log2Fold 2.19), and *aoxA1* (Log2Fold 2.90) (**Figure 2A** and **Supplementary Table 8**). However, at the 75-minute time point, additional GO term categories such as light induced genes (*brlA* μORF, *brlA, prtA*, and AN0045) are up-regulated. The last time point had nine light-induced DE up-regulated genes (*brlA, prtA*, AN0045, AN0693, AN1893, AN3304, AN3872, AN5015, AN8641

**Figure 2:**
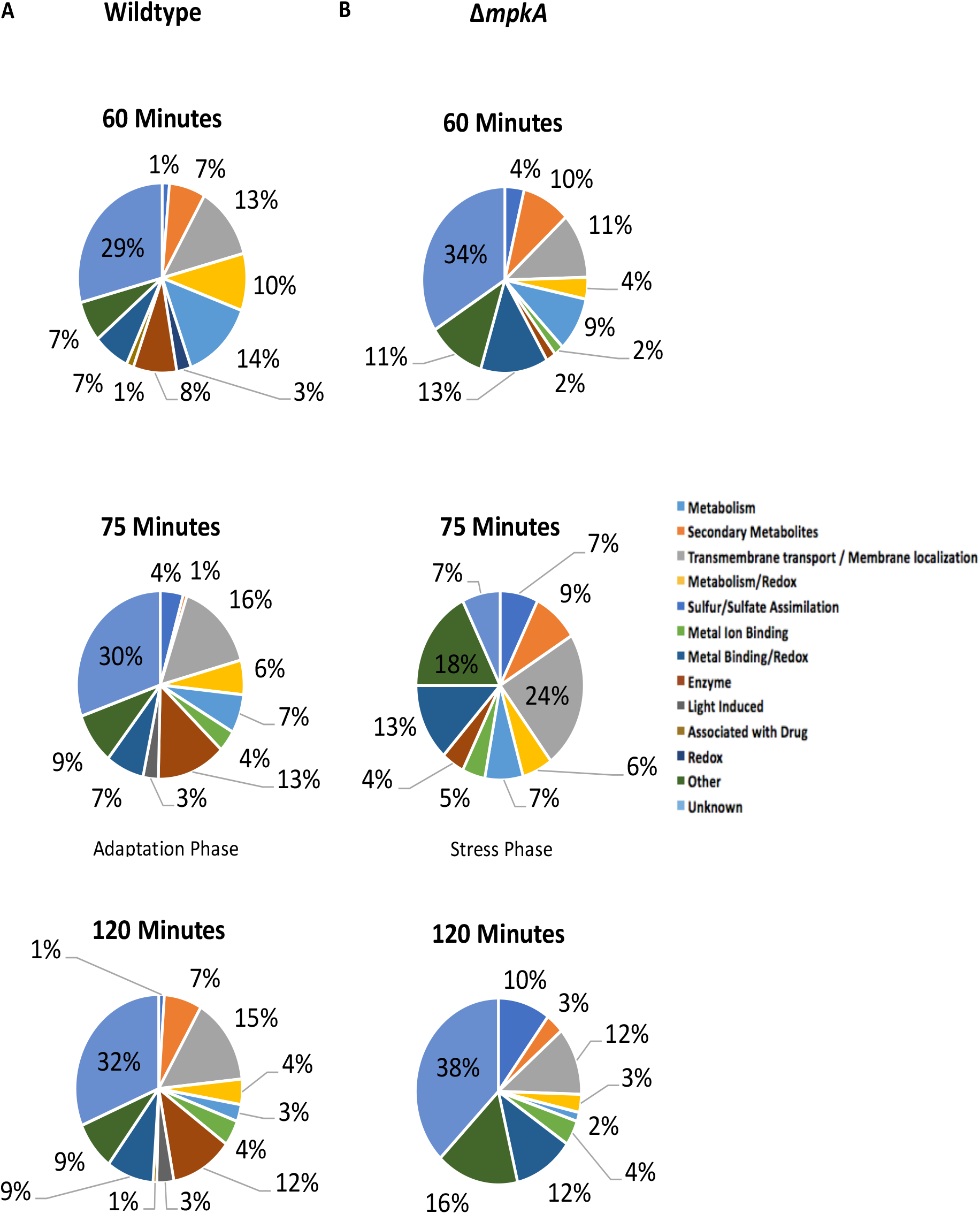
Gene Expression Shifts at the 75 Minute Time Point in Wildtype. **A)** Up-regulated wildtype gene expression at time points 60, 75, and 90-minutes **B)** Up-regulated Δ*mpkA* gene expression at time points 60, 75, and 90-minutes. GO Terms from *Aspergillus* Database.

(**Figure 2A** and **Supplementary Table 8**). Also, there is a known connection with secondary metabolism and light induced genes, which is particularly evident at the 75-minute time point and beyond. Examples include *pkdA* (Log2Fold 4.3), *catA* (Log2Fold 2.49), *amyF* (Log2Fold 2.5), AN9314 (Log2Fold 5.79), and AN2116 (Log2Fold 6.34)) (**Figure 2A** and **Supplementary Table 8**) [21]. Notably, the distribution of DE up-regulated GO terms for the Δ*mpkA* mutant does not display this apparent shift, as there are no light-induced up-regulated in any of the time points of the Δ*mpkA* strain (**Figure 2A** and **Supplementary Table 8**).

### 3.4 Light Induced Gene Induction

Further investigation of genes that exhibit increased expression upon exposure to micafungin in wildtype revealed the induction of *brlA* (AN0973) and *brlA* μORF (AN0974) at 75, 90, and 120-minutes (**Supplementary Table 4**). *brlA* was up-regulated at Log2Fold change of 2.70, 3.11, 3.50 at time points 75, 90, 120-minutes, respectively (**Figure 3A** and **Supplementary Table 6**). *brlA* μORF was up-regulated with Log2Fold changes of 3.5, 3.2, 3.1 at time points 75, 90, 120-minutes, respectively (**Figure 3A** and **Supplementary Table 4**). qRT-PCR results at 75-minutes after micafungin exposure confirmed the significant up-regulation of *brlA* (7.8 log fold change) and *brlA* μORF (12.09 log fold change) in the presence of micafungin in wildtype but not in the *mpkA* knockout (*brlA* 0.91 log fold change; *brlA* μORF 1.52 log fold change) (**Figure 3B**).

**Figure 3:**
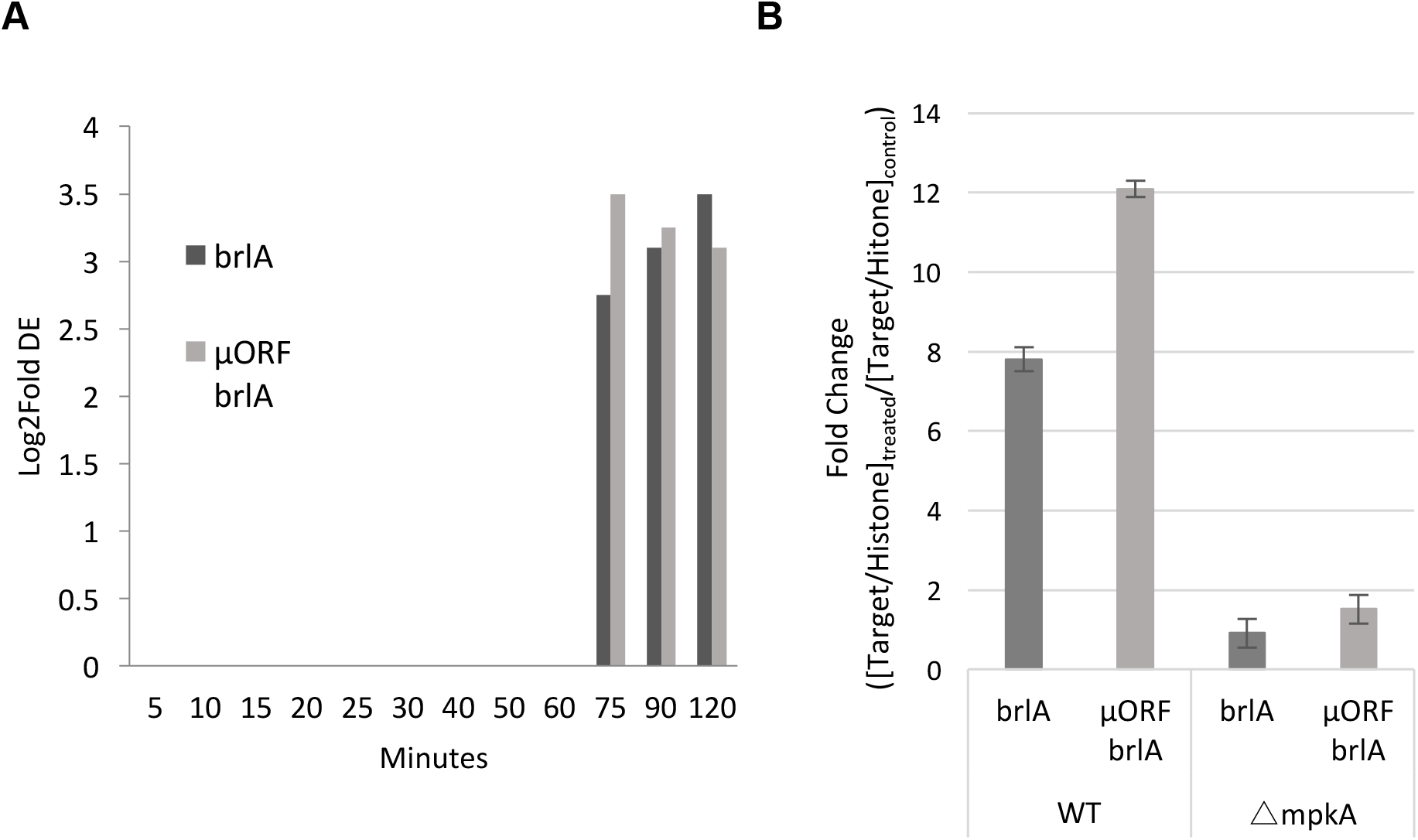
*brlA* Up-regulated in Wild Type but not Δ*mpkA*. **A)** RNA-Sequencing data showing *brlA* and *brlA* μORF up-regulated in wild type starting the adaptation phase. 3 biological replicates. Significance DE = Log2Fold >2.0. **B)** qRT-PCR at the last time point, *brlA* and *brlA* μORF are up-regulated in wild type but not in Δ*mpkA*. Three biological replicates.

The induction of *brlA* expression in wildtype hyphae exposed to cell wall damage was unexpected. *brlA* encodes a C_2_H_2_ zinc finger transcription factor that regulates conidiophore development [15]. However, asexual structures typically do not develop in aerated shaking flask cultures as were used in this study (i.e., hyphae should only grow in a vegetative state) [22]. In the well-characterized regulatory pathway that controls asexual development in *A. nidulans*, upstream regulators activate *brlA* expression, which then results in the induced expression of the downstream transcription factors *abaA* and *wetA* [15]. Also, there was no significant increase of transcript expression for *abaA* (AN0422) (Log2Fold 0.329 at time point 75-minutes) or *wetA* (AN1937) (Log2Fold 0.290 at time point 75-mintues) upon exposure to micafungin (**Supplemental Table 4** and **Supplemental Table 5**), nor were any know upstream activators affected (data not shown). These results suggest that the observed increased expression of *brlA* does not reflect the full induction of the canonical asexual development regulatory pathway.

### 3.5 Hyphal Response to Micafungin exposure

To investigate the possibility that expression of brlA might correlate with specific morphological changes. We exposed wildtype and Δ*mpkA* hyphae to 0 ng/ml (untreated control), 0.01 ng/ml, 0.1 ng/ml, and 1.0 ng/ml of micafungin in YGV media after they had already been growing for 11 hours on cover slips at 28°C. We let the strains grow under these same conditions for an additional three hours post-treatment. Whereas the untreated control strains exhibited no morphological alterations (**Figure 4**), wildtype hyphae exposed to micafungin possessed bulges at hyphal tips and lateral branching points regardless of micafungin dosage (**Figure 4**).

**Figure 4:**
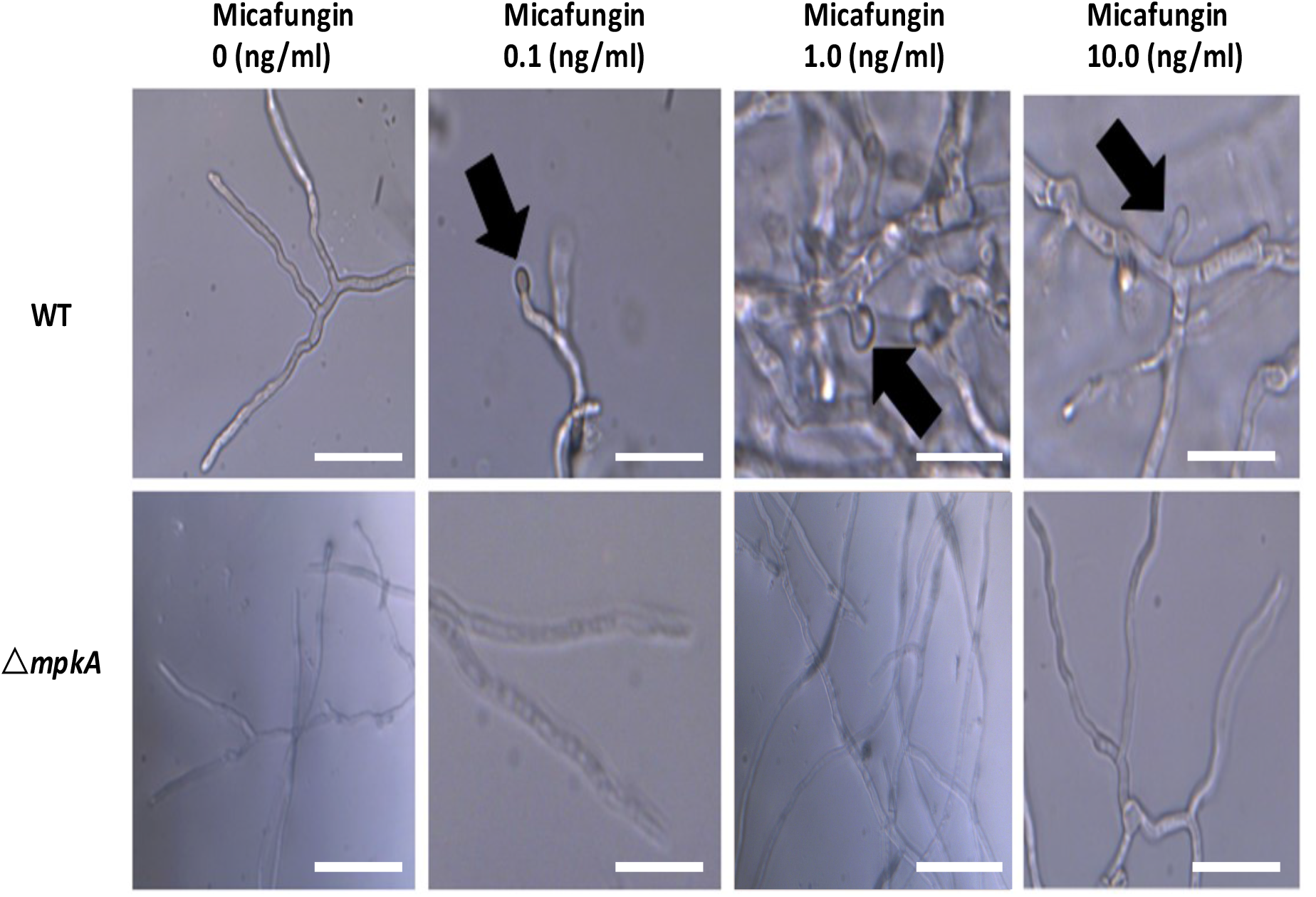
Hyphal morphology following exposure to Micafungin. The morphology of wildtype and Δ*mpkA* hyphae that were untreated was normal. Wildtype hyphae display bulges at hyphal tips and lateral branches when exposed to micafungin at each tested dosage. No morphological differences in Δ*mpkA* hyphae following micafungin exposure. 60X magnification. Scale bar is 25µm. Pictures taken on Olympus BX51 microscope.

The bulging phenotype of wildtype hyphae exposed to micafungin resembled a pattern of attenuated spore development in filamentous fungi that is known as microcycle conidiation [9]. This phenomenon has been observed in *Aspergilli* exposed to poor growth substrates such as indoor construction materials or hospital air filters [23]. To test the idea that microcycle conidiation is a response to micafungin exposure, we repeated the previous experiment but extended the post-dosage incubation period to allow for spore formation. Accordingly, wildtype and Δ*mpkA* hyphae were grown and treated with micafungin as described above, with the difference that hyphae were incubated for nine hours instead of three prior to observation. In untreated control hyphae, no morphological alterations were observed (**Figure 5**). However, wildtype hyphae exposed to 0.1ng/ml of micafungin exhibited development of spores at hyphal tips (**Figure 5**). Cover slips were stained with calcofluor to confirm the development of spores at hyphal tips (**Figure 5**). Unlike wildtype hyphae, the Δ*mpkA* strain did not exhibit development of spores post 9 hours of micafungin exposure (**Figure 5**).

**Figure 5:**
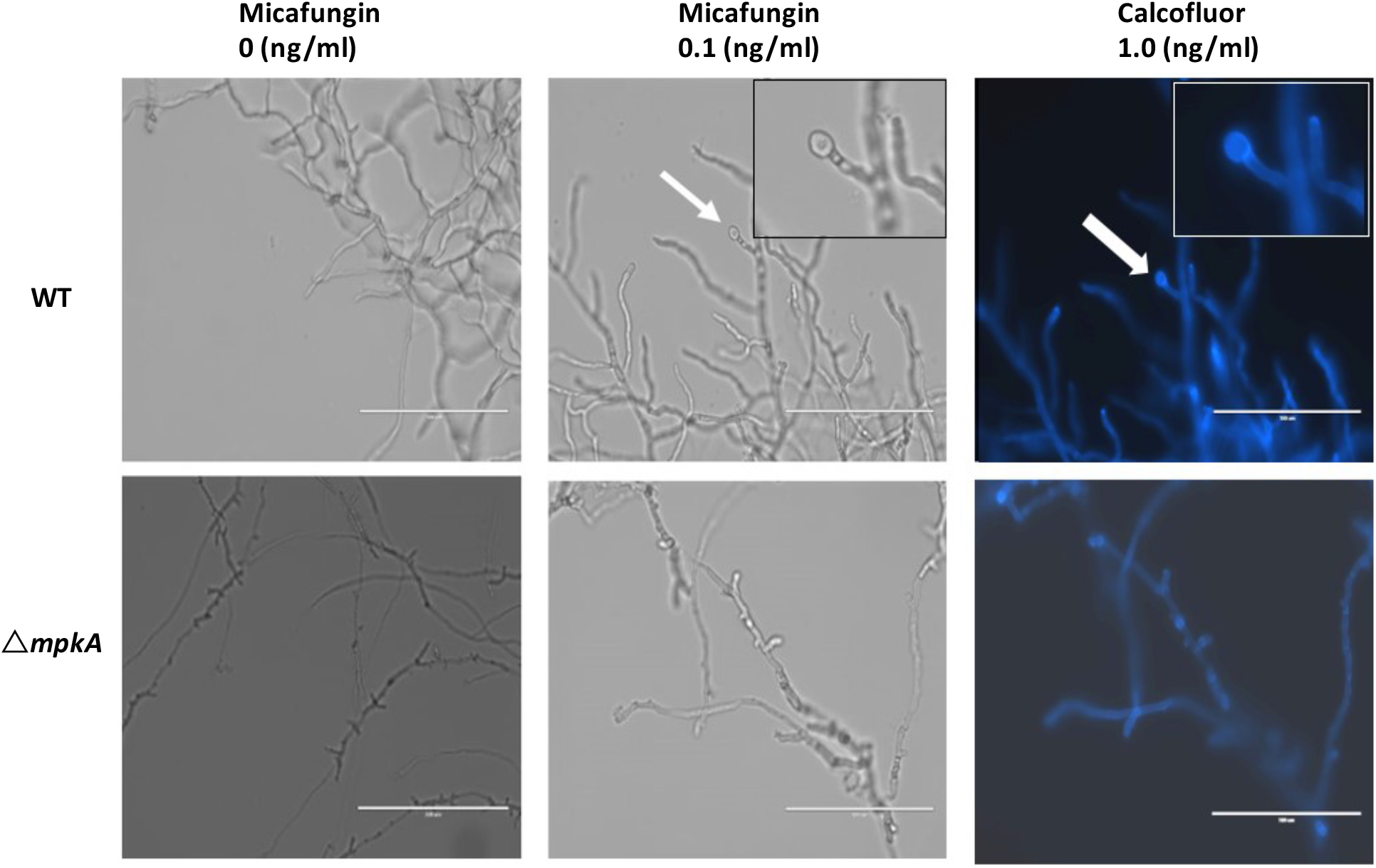
Microcycle Conidiation Spore Development Post Micafungin Exposure. Microcycle conidiation develops spore at hyphal tips in wildtype post nine hours exposure to micafungin. No spore development in Δ*mpkA*. 40X magnification. Scale bar is 100µm. Pictures taken on EVOS FL microscope.

### 3.6 Similar spore morphologies upon Induced brlA Expression and Post Micafungin Exposure

We sought to determine whether the spores generated by micafungin-induced microcycle conidiation are similar to those that result from established conditions known to trigger this process. Previously, it was demonstrated that forced activation of *brlA* expression results in production of a single spore at hyphal tips and branching sites [15,16]. As this is effectively a form of microcycle conidiation, we compared spores produced upon forced induction of *brlA* to those generated as a result of micafungin exposure. The *alcA::brlA* strain was grown on cover slips for three hours in YGV media at 28C then shifted to MV 100mM L-Threonine to grow for an additional 3 hours to induce *brlA* expression. The wildtype strain was grown and exposed to micafungin for nine hours by the same methods as described above. Both strains were viewed under the microscope. The *alcA::brlA* strain developed spores at the hyphal tips, as did the wildtype strain exposed to micafungin (**Figure 6**). The hyphae were stained with calcofluor at 1.0ng/ml to clearly show the development of spores (**Figure 6)**. There were similarities to the morphology of spores produced by both strains being the spore size similarities and calcofluor stain intensity. These similarities suggest that BrlA-induced microcycle conidiation is a key response to cell wall damage caused by micafungin in *A. nidulans* hyphae.

**Figure 6:**
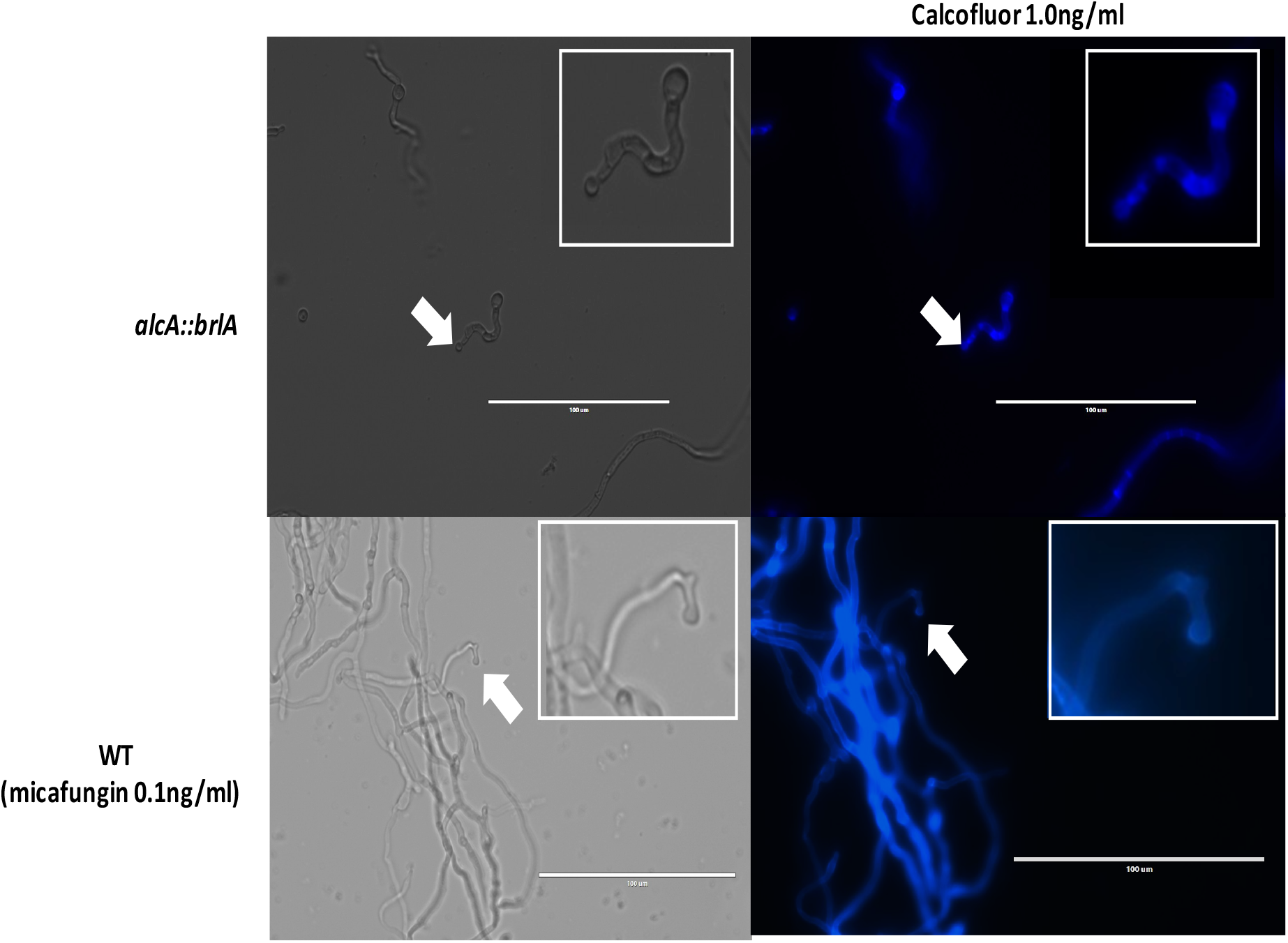
Similar Microcycle Spore Generation. Similar spore phenotypes in up-regulated *brlA* strain after three hours promoter induction and wildtype nine hours post micafungin exposure. 40X magnification. Scale bar is 100µm. Pictures taken on EVOS FL microscope.

## 4. Discussion

The purpose of this study was to investigate the dynamic transcriptional response of *A. nidulans* to the antifungal drug micafungin, which compromises cell wall integrity through the depletion of β-1,3-glucans. We compared the response of wildtype hyphae to that of an Δ*mpkA* deletion mutant, in which the terminal kinase of the CWIS pathway has been disabled. In a prior study, our characterization of the dynamic phosphoproteomic and transcriptional response of wildtype hyphae revealed candidate signaling and morphogenetic pathways that promote repair of damaged cell walls and the maintenance of hyphal integrity [6]. Here, we show that a significant transcriptional response still occurs in the absence of MpkA. Nevertheless, MpkA is required for expression of a set of genes that seemingly support the long-term survival of *A. nidulans* in the face of cell wall damage. Unexpectedly, amongst these genes is the master regulator of asexual development, *brlA*. Further investigation suggested that expression of BrlA enables microcycle conidiation, which might represent a heretofore unconsidered strategy used by fungi to resist the impacts of cell wall damage.

The distinct features of the *A. nidulans* CWIS pathway compared to the established yeast paradigm were first noted by Fujioka et al (2007), who observed that the response in filamentous fungi includes a significant MpkA-independent component [3]. They found that expression of several cell wall-related genes, particularly those involved in β-1,3-glucan and chitin synthesis, did not require functional MpkA. Our results further underscore the complexity of the *A. nidulans* response, as we observed a significant transcriptional response to cell wall damage despite the absence of MpkA (∼1000 genes compared to 1600 in wildtype). However, two features of the response in Δ*mpkA* mutants were notable. First, the timing of significant up-regulation of gene expression appears to be delayed by about ten minutes in Δ*mpkA* mutants relative to wildtype. Thus, a key feature of the MpkA-dependent component of the CWIS might be temporal, which would presumably ensure a rapid “all hands-on deck” response to repair the damaged cell wall. Second, certain classes of genes whose expression is up-regulated do in fact appear to be dependent upon the presence of functional MpkA. This includes sets of genes implicated in responses to light and in secondary metabolism.

Other than the observation that their expression is induced by light, the function of most of the genes that fall into the former class remains unknown with the exception of *brlA* (see below) and *conJ* (AN5015) [24, 25]. Amongst those genes that fall into the latter class, a small set of gene clusters potentially involved in secondary metabolite biosynthesis exhibited MpkA-dependent up-regulation in response to micafungin. On the other hand, another equally small set displayed MpkA-independent up-regulation.

Other than underscoring the complexity of the transcriptional response to micafungin, the implications of the observed up-regulation of these gene clusters awaits their functional characterization. Of note, several additional gene clusters, including some encoding known polyketide synthases or non-ribosomal peptide synthases showed dynamic MpkA-dependent expression even though they were not induced by micafungin [26]. This presumably reflects the established role of MpkA in the regulation of secondary metabolism though the underlying mechanism remains to be determined [18, 27].

Microcycle conidiation has been observed in a variety of filamentous fungi, and generally appears to serve as a “fail-safe” mechanism that enables fungi to efficiently divert resources from cellular growth to sporulation under adverse conditions [10, 9]. The types of stress reported to trigger microcycle conidiation include nutrient depletion, high temperatures, altered pH, and high cell density [9]. The morphological steps that underlie microcycle conidiation vary across fungi, and relatively little is known about the precise mechanisms that subvert growth and cause sporulation. In *A. nidulans*, it has been previously shown that forced expression of *brlA* under conditions that normally do not promote sporulation results in the cessation of growth and the production of terminal conidia [15]. Although somewhat artificial, this form of microcycle conidiation requires a functional Cdc42 GTPase module [16]. Here, we show that micafungin-induced expression of *brlA* leads to an analogous form of microcycle conidiation. The involvement of developmental regulators such as *veA* and *wetA* in the regulation of microcycle conidiation has been noted in other fungi [11, 28]. These results support the notion that the regulatory factors and pathways that control normal forms of asexual reproduction in fungi also support microcycle conidiation. Our results further highlight the importance of microcycle conidiation as a survival strategy for fungi that encounter adverse environmental conditions. We propose that the CWIS is a multi-faceted response that is primarily dedicated to maintaining the integrity of growing hyphae through the repair of damaged cell walls. The existence of additional outputs such as microcycle conidiation conceivably provides alternative survival options should the damage be persistent or irreparable.

The echinocandins are an efficacious class of anti-fungal drugs that include micafungin [29]. However, the usefulness of the echinocandins is threatened by the emergence of resistance in fungi such as *Candida* species, and the genetic analysis of resistant isolates shows that they often harbor mutations in genes other than the known target *FKS1* [30, 31]. These other genes appear to primarily control processes that compensate form reduced cell wall β-glucans. Our results imply that microcycle conidiation represents another compensatory mechanism that might contribute to echinocandin resistance, and suggest that further mechanistic study of this survival strategy might facilitate discovery of new approaches for countering resistance.

## 5. Conclusions

Filamentous fungi deal with stress at the cell wall through the CWIS pathway [3]. This stress can be caused by many factors including developing a weak cell wall due to exposure of an antifungal [4]. The understanding of how the fungi responds to this stress can lead to a better understanding of fungal resistance to antifungals and can help improve the understanding of the different categories of antifungals. This study showed that there are two phases in response to an echinocandin antifungal in *Aspergllus nidulans*, and certain genes can be induced to regulate the switch between the phases. This phase switch is evolutionarily beneficial to the fungi as it can preserve the genomic DNA in an efficient and timely manner. The knowledge gained in this study can help the development of antifungal drugs to expand the types of antifungals offered to treat infected patients by developing a deeper understanding of the echinocandin drug category.

## Supplementary Materials

Figure S1: Shared DE Genes, Table S1: Strains used in this Study, Table S2: NCBI SRAs, Table S3: qRT-PRC Primers, Table S4: WT RNA-Seq Data, Table S5: KO RNA-Seq Data, Table S6: RNA-Sequencing DE Genes, Table S7: Gene Clusters, Table S8: Up-regulated GO Terms for RNA-Seq

## Funding

The research conducted in this publication was funded by the NASA Grant (80NSSC17K0737) and by the NSF Grant (MCB 1601935).

## Author Contributions

S.R., C.C., W.R., M.R.M., and S.H. designed research; C.C. and S.R. performed research; S.R. analyzed data; S.R., W.R., and S.H. wrote the paper. All authors have read and agreed to the published version of the manuscript.

## Acknowledgments

We acknowledge Jenna Van Bosch from the University of Nebraska-Lincoln for preformed research.

## Institutional Review Board Statement

NA

## Informed Consent Statement

NA

## Data Availability Statement

RNA-Sequencing data deposited to NCBI and BioProject numbers are: PRJNA562694 and PRJNA727274. Additional information needed contact the corresponding author.

## Conflicts of Interest

Authors declare no competing interests.

## Notes

### Competing Interest Statement

The authors have declared no competing interest.

