## Supplementary material for "Micafungin-induced Cell Wall Damage Stimulates Microcycle Conidiation in *Aspergillus nidulans*": SF1

A

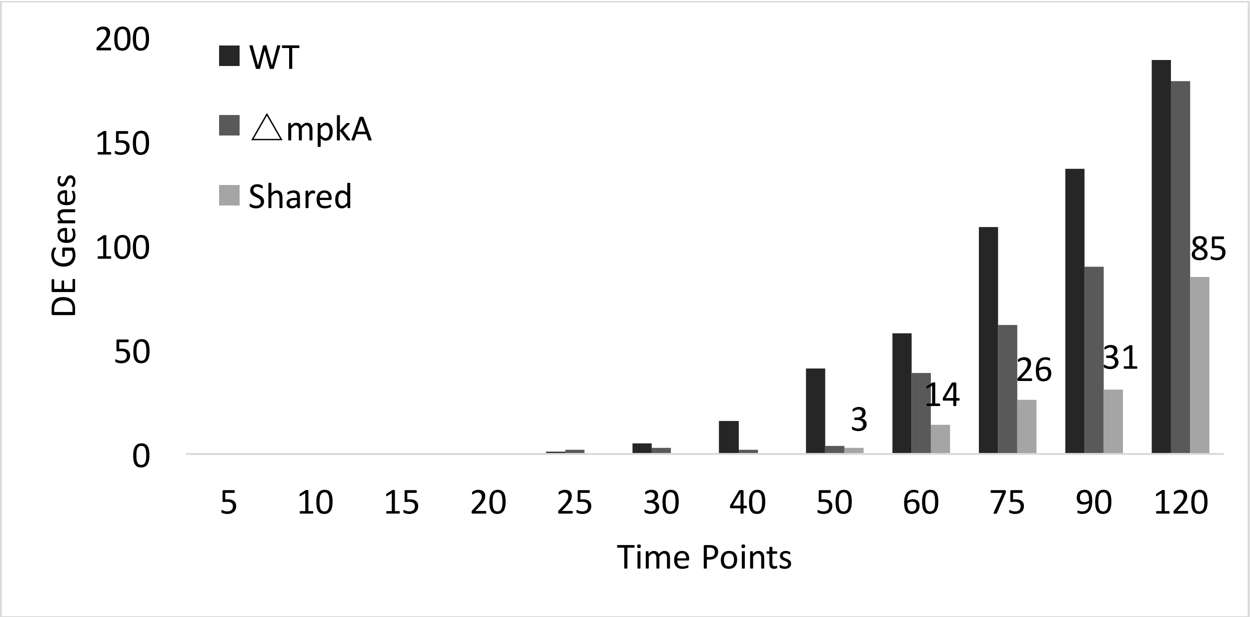


B

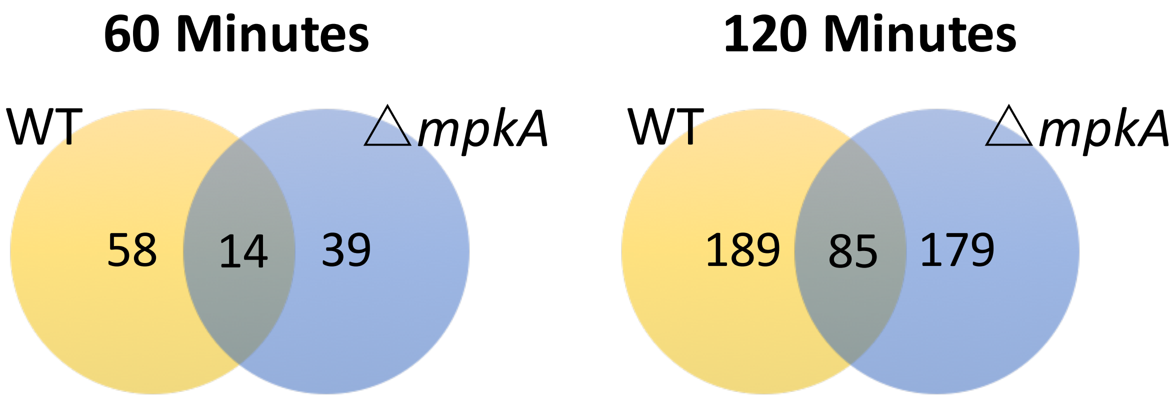


**<u>Supplemental Figure 1.</u> A** Up-regulated DE genes that were shared by both wildtype and Δ*mpka* through all the time points. The first shared gene occurs at 50 minutes. **B** Time point 60-minutes there are 14 shared DE genes and at time point 120 minutes there are 85 shared
